# *In vivo* characterization of a secologanin transporter from *Catharanthus roseus*

**DOI:** 10.1101/2023.08.01.551495

**Authors:** Fanfan Li, Mohammadamin Shahsavarani, Cody-Jordan Handy-Hart, Victoria Montgomery, Robin N. Beech, Lan Liu, Stéphane Bayen, Yang Qu, Vincenzo De Luca, Mehran Dastmalchi

## Abstract

Monoterpenoid indole alkaloid (MIA) biosynthesis in *Catharanthus roseus* is a paragon of the spatiotemporal complexity achievable by plant specialized metabolism. Spanning a range of tissues, four cell types, and five cellular organelles, MIA metabolism is intricately regulated and organized. This high degree of metabolic differentiation requires inter-cellular and organellar transport, which remains understudied. Here, we have fully characterized a vacuolar importer of secologanin belonging to the multidrug and toxic compound extrusion (MATE) family, named CrMATE1/SLTr. Phylogenetic analyses of MATEs suggested a role in alkaloid transport for CrMATE1, and *in planta* silencing in two varieties of *C. roseus* resulted in a shift in the secoiridoid and MIA profiles. Subcellular localization of CrMATE1 confirmed tonoplast localization. A full panel of *in vivo* biochemical characterization using the *Xenopus laevis* oocyte expression system was used to determine substrate range, directionality, and rate. We can confirm that CrMATE1 is a vacuolar importer of secologanin, rapidly transporting 1 mM of secologanin within 25 min. Notably, the absence of CrMATE1 leads to a transport bottleneck, resulting in the conversion of secologanin to its reduced form, secologanol, both *in planta* and in the *X. laevis* system. The unique substrate-specific activity of CrMATE1 showcases the utility of transporters as gatekeepers of metabolic flux, mediating the balance between anti-herbivory potency and cell homeostasis *in planta*. MIA and secoiridoid transporters could also be deployed in heterologous systems to guide biosynthetic pathways and improve titers of valuable and life-saving MIAs.

**SIGNIFICANCE:** We have fully characterized CrMATE1, a multidrug and toxic compound extrusion (MATE) family transporter in *Catharanthus roseus,* as a vacuolar importer of secologanin. The translocation of secologanin into the vacuole is necessary for the first committed step of monoterpenoid indole alkaloid (MIA) biosynthesis.

## 1. INTRODUCTION

The patterning of specialized chemical profiles throughout the plant body is complex, expectedly, and an indication of resource allocation and metabolic differentiation (St-Pierre *et al*., 1999; Li *et al*., 2023). It is also a means for dedicated storage (Hagel *et al*., 2008) and rapid deployment or activation of potentially toxic components (Morant *et al*., 2008; Guirimand *et al*., 2010; Vogel *et al*., 2010), which requires inter-/intra-cellular transport. The movement of specialized metabolites can occur at a proximal or distal level, at a subcellular resolution or encompassing the full range of plant organs. In both cases, transporters are often required to facilitate movement across semi-permeable membranes, such as the vacuolar (tonoplast) or cellular/plasma membranes.

An intricate picture of compartmentalization has been developing in the study of monoterpenoid indole alkaloids (MIAs) in the medicinal and horticultural plant *Catharanthus roseus* (L.) G. Don (St-Pierre *et al*., 1999; Courdavault *et al*., 2014; Kulagina *et al*., 2022; Sun *et al*., 2022; Li *et al*., 2023). Many transporters have been identified for the multi-step biosynthetic pathway culminating in the anticancer bisindole alkaloids vinblastine and vincristine (Figure 1). A series of Nitrate Peptide Family (NPF) transporters (CrNPF2.4-2.6) have been characterized with H^+^-coupled import of the secologanin biosynthetic intermediates (deoxyloganic acid, loganic acid, loganin and secologanin) into the cell, across the plasma membrane (Larsen *et al*., 2017). These transporters, or a subset, could putatively transport secoiridoids from the roots to the leaf epidermis, as the inter-organ accumulation of secologanin has been suggested (Kidd *et al*., 2019). Conversely, CrNPF2.9 is a tonoplast-localized exporter of strictosidine, the reaction product of secologanin and tryptamine condensation, which presumably occurs in the leaf epidermis, providing the first committed intermediate of the MIA pathway (Payne *et al*., 2017).

**Figure 1.**
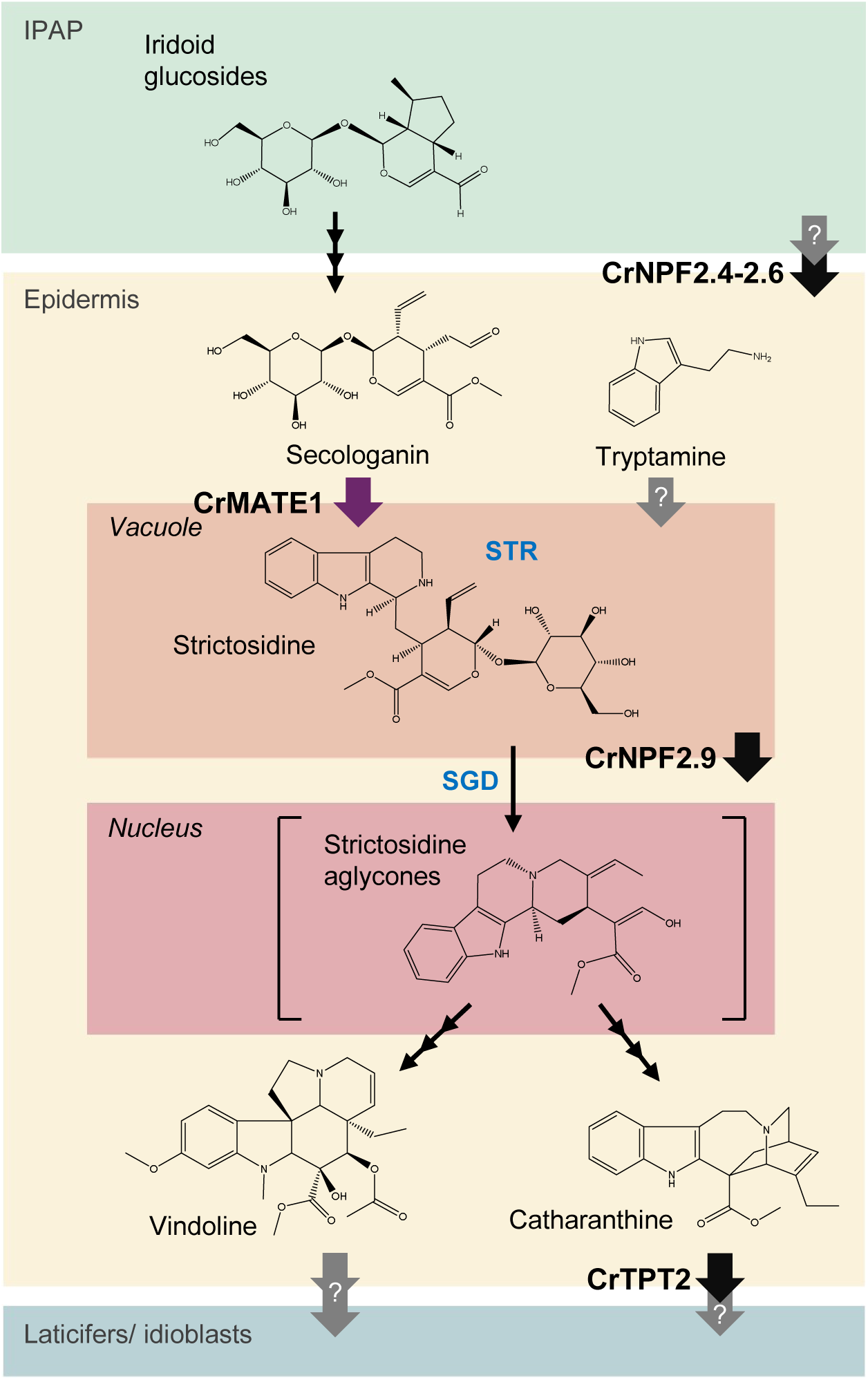
Compartmentalization of resolved and unknown transport steps in the *C. roseus* MIA pathway. Biosynthesis of iridoid glucosides derived from the MEP pathway occurs in the internal phloem-associated parenchyma (IPAP) and they are transported out via an unknown mechanism. Secoiridoids are moved into epidermal cells via CrNPF2.4-2.6 (Larsen *et al*., 2017), where secologanin is imported to the vacuole by CrMATE1 (purple arrow). An unknown transporter concurrently imports tryptamine. In the vacuole, condensation of the two molecules by strictosidine synthase (STR) forms the central MIA intermediate, strictosidine, which is exported into the cytosol by CrNPF2.9 (Payne *et al*., 2017). Strictosidine-β-D-glucosidase (SGD) guides its substrate into the nucleus, where strictosidine aglycones accumulate. A series of enzymatic steps leading to catharanthine follows, where it is finally exported via CrTPT2 (Yu and De Luca, 2013), accumulating in the laticifers/idioblasts, along with vindoline, mediated by unknown mechanisms. Uncharacterized transport steps are shown with grey arrows. Multiple reaction steps are shown with three conjoined arrows.

Further downstream, an ATP Binding Cassette (ABC)-type transporter has been identified as a catharanthine plasma membrane exporter (CrTPT2) that is expressed predominantly in the epidermis of young leaves (Yu and De Luca, 2013). This class of TPT2 transporters, belonging to the pleiotropic drug resistance (PDR) subfamily, appears to be conserved across the Apocynaceae family, with an ortholog from the related species *Vinca minor* (VmTPT2/VmABCG1) characterized with a similar function (Demessie *et al*., 2017). These transporters appear to play a role in the accumulation of catharanthine and vincamine, respectively, with several reports suggesting that the idioblasts and laticifers are likely the final storage site (Yamamoto *et al*., 2016; Sun *et al*., 2022; Guedes *et al*., 2023; Li *et al*., 2023).

A recent report has suggested that a member of the multidrug and toxic compound extrusion (MATE) protein, SLTr (hereafter referred to as CrMATE1), transports secologanin across the tonoplast, where it would be available to strictosidine synthase (STR) (Li *et al*., 2023). Interestingly, CrMATE1 forms a cluster on chromosome 3 with STR and tryptophan decarboxylase (TDC), which catalyze consecutive reactions (Sun *et al*., 2022; Li *et al*., 2023). The role of this MATE was inferred based on virus-induced gene silencing (VIGS) of *CrMATE1* in young *C. roseus* leaves. The authors reported no change in secologanin levels but an increase in its reduced form, secologanol. It was reasoned that secologanol build-up was a by-product of secologanin remaining aberrantly in the cytosol. However, the authors could not conduct *in vitro* or *in vivo* transport assays due to the recalcitrance of the gene to heterologous expression. In parallel, our group first became interested in this MATE protein after the publication of the *C. roseus* genome by Kellner *et al*. (2015). We further pursued characterization of the transporter when it was identified (CL1653Contig1, EST=2) in a leaf epidermis (LE)-enriched transcriptome (Murata *et al*., 2008). Evidence from single-cell and idioblast-targeted transcriptomics has supported the theory that this transporter, along with several others, is enriched in epidermal, laticifer, and internal phloem-associated parenchyma (IPAP) cell types. (Sun *et al*., 2022; Guedes *et al*., 2023; Li *et al*., 2023). However, our main challenge lay in provisioning an efficient heterologous platform for the direct characterization of CrMATE1.

Our analyses recapitulate the VIGS-based silencing of *CrMATE1*, which we conducted independently in two different *C. roseus* backgrounds, “Little Delicata (LD)” and “Pacifica White (PW).” Silencing of *CrMATE1* in both backgrounds led to reduced secologanin and a drastic accumulation of secologanol, possibly due to spontaneous or endogenously-catalyzed reduction of the aldehyde. We confirmed the tonoplast localization of CrMATE1 by visualizing the eGFP fusion protein in *Nicotiana benthamiana*. Finally, we biochemically characterized CrMATE1 using *Xenopus laevis* oocytes as a proxy for the plant vacuole. Using this expression platform, we confirmed that CrMATE1 is exclusively a vacuolar importer of secologanin, having tested other secoiridoid intermediates. Overall, these results establish the localization, directionality, substrate, and rate of this MATE secologanin transporter, improving our understanding of the mechanics of MIA spatiotemporal distribution.

## 2. RESULTS

### 2.1 Phylogenetic analysis of MATEs

We compiled MATEs from plant and prokaryotic sources, with representatives from the major subfamilies (groups A-F) and constructed a maximum likelihood tree (Suppl. Figure 1). In this phylogenetic tree, plant MATEs formed four clades: A, B, C, and F. Group A includes CrMATE1 and many known transporters of alkaloids and flavonoids (Gani *et al*., 2021). CrMATE1 appears closely related to NtMATE1 and 2, which have been characterized as transporters of nicotine in *Nicotiana tabacum* (Shoji *et al*., 2009).

MATE proteins have been described as forming a bilobate V-shape structure, with pseudo-symmetrical assembly of the N- and C-lobes (Tanaka *et al*., 2021). While MATE proteins belonging to groups B and C have charged, hydrophilic residues in the V-shaped pocket (E136, E379, E384, and R508), group A members have hydrophobic or uncharged substitutions at some of these residues (Suppl. Figure 2). Here we show that CrMATE1 and NtMATE1 share the same substitutions for two out of the four hydrophilic residues conserved in groups B and C. Hydrophilic residues E136, K275, and E379 are conserved in Group A, while E384Q and R508V are replaced with uncharged amino acids for both CrMATE1 and NtMATE1.

### 2.2 Silencing of *CrMATE1* reduces secologanin content

Virus-induced silencing (VIGS) was conducted to suppress the expression of *CrMATE1* in *C. roseus* leaves, referred to hereafter as *VIGS-MATE*, compared to empty vector (*EV*)-infiltrated plants. Silencing was carried out in two *C. roseus* varieties, “LD” and “PW.” A 255 bp region was selected for silencing (Suppl. Figure 3A), and qPCR was used to confirm the silencing of *CrMATE1*. Expression of *CrMATE1* was reduced by 73 and 80% in LD and PW backgrounds, respectively (Figures 2A, C).

**Figure 2.**
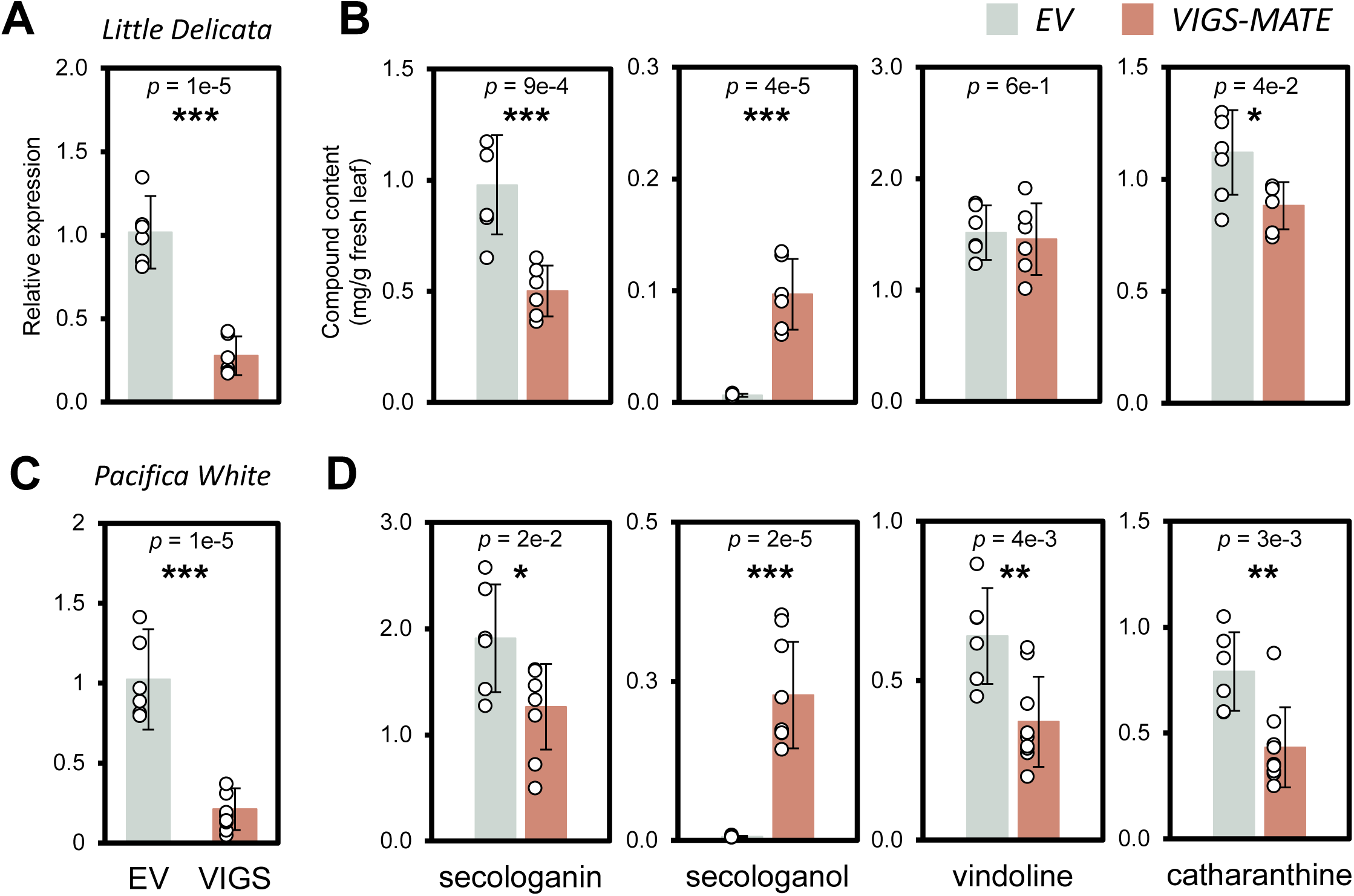
Silencing *CrMATE1* leads to an increase in secologanol. **A)** In the background “Little Delicata (LD),” *CrMATE1* transcripts were reduced by 73% (*VIGS-MATE*) compared to the empty vector (*EV*) controls. **B)** Chemical profile of LD *VIGS-MATE* reveals a 50% reduction in secologanin, and a 16-fold increase in its reduced form, secologanol, compared to the *EV*. Vindoline was not affected and catharanthine was significantly reduced. **C)** In “Pacifica White (PW),” *CrMATE1* expression was suppressed by 80%. **D)** Chemical profile of PW *VIGS-MATE* shows a significant reduction in secologanin and a 38-fold increase in its reduced form, secologanol, compared to the *EV*. Vindoline and catharanthine were significantly reduced by 42% and 46%, respectively. Data represents the mean ± S.D. of six biological replicates, except for PW *VIGS-MATE* (nine biological replicates). Data points are shown. One, two, and three asterisk(s) signify *p* < 0.05, *p* < 0.01, and *p* < 0.001, respectively.

Silencing of *CrMATE1* in both backgrounds led to a significant reduction in secologanin (Figures 2B, D). In PW, a reduction was also seen in downstream MIAs, vindoline and catharanthine (Figure 2D). However, in LD only catharanthine was significantly reduced (Figure 2B). Notably, there was a drastic accumulation of secologanol in both varieties, a reduced form of secologanin. Present only as a minor peak in the *EV* plants, secologanol was identified by matching MS/MS spectra with a chemically produced standard (Suppl. Figure 3B, C). Secologanol increased in silenced plants by 16 to 38-fold in LD and PW backgrounds, respectively.

### 2.3 CrMATE1 is localized to the tonoplast

Using fluorescence confocal microscopy, eGFP-CrMATE1 was shown to localize to the tonoplast. CrMATE1 displayed the hallmarks of tonoplast localization, including circular extensions or ‘bulbs’ into the lumen of the vacuole and transvacuolar strands, seen streaming across the cellular confines (Figures 3A, D). Furthermore, a clear separation of fluorescence between adjacent cells is apparent, indicative of localization to the tonoplast, as compared to the plasma membrane (Figure 3C). These observations were further confirmed by co-expressing eGFP-CrMATE1 with a vacuolar marker fused to RFP (Suppl. Figure 4) (Nelson *et al*., 2007). In conjunction, *CrMATE1* expression in *C. roseus* was quantified in several tissue and developmental samples by qPCR, including stem, root, flower, and leaf pairs (1-3) (Suppl. Figure 5). Expression was ubiquitous throughout sampled tissues but with slightly higher expression in the leaf.

**Figure 3.**
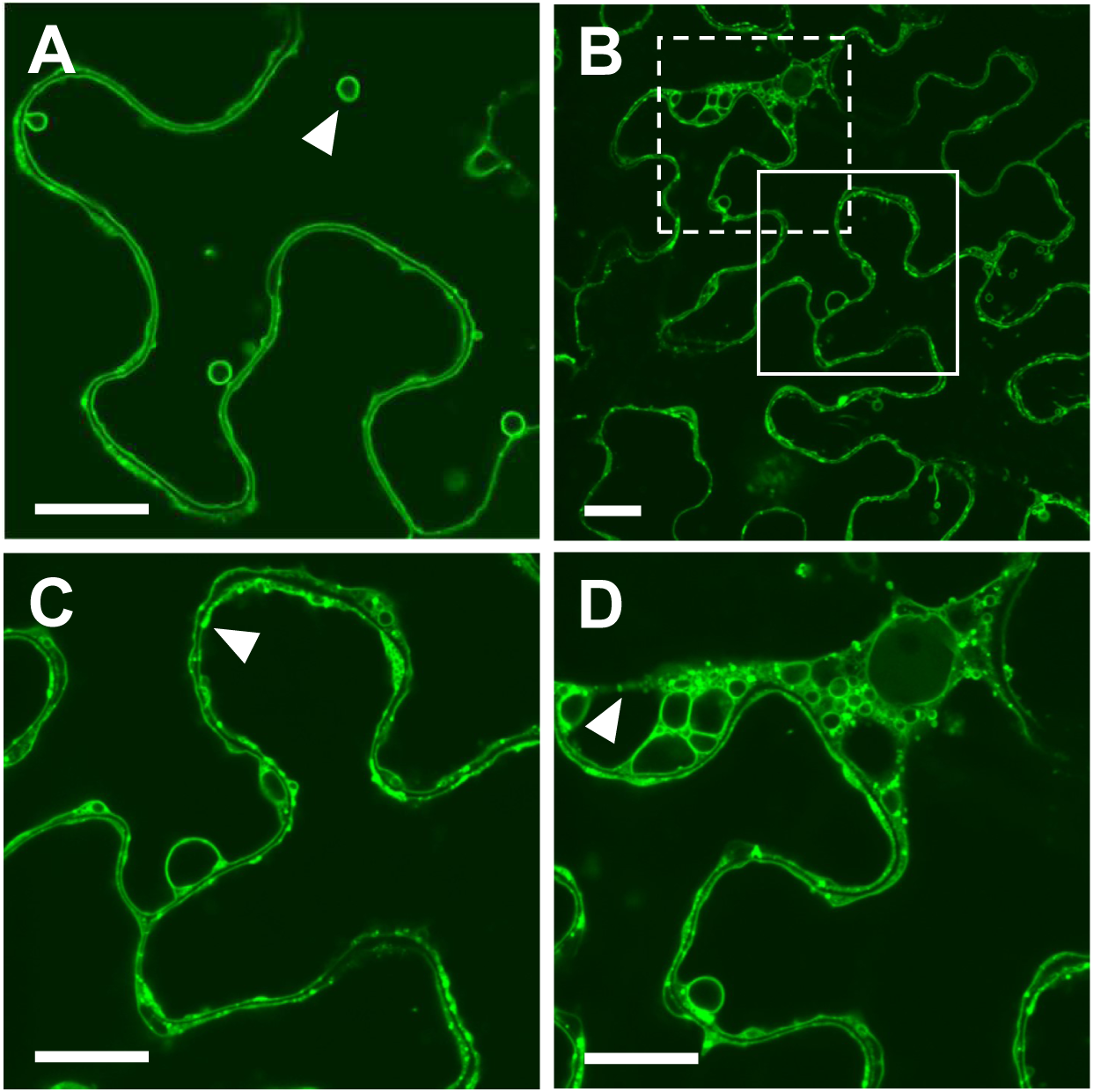
CrMATE1 localizes to the tonoplast. Fluorescent reporter eGFP was fused to the N-terminus of CrMATE1 and transiently expressed in *N. benthamiana* leaf epidermal cells: **A)** CrMATE1 localized to the tonoplast, with circular extensions ‘bulbs’ into the lumen of the vacuole; **B)** CrMATE1 localization to the tonoplast with higher magnification to reveal **C)** clear tonoplast separation between adjacent cells, and **D)** transvacuolar strands, streaming across the cell. Features are highlighted with white arrowheads. Magnified areas from panel B are denoted by white and dashed boxes, and are shown in panels C and D, respectively. Subcellular localization was determined by surveying more than 50 cells in each of the three biological replicates. Scale bars indicate 20 µm.

### 2.4 Biochemical characterization of secologanin transport

The results of *in planta* silencing and subcellular localization suggested that CrMATE1 was a vacuolar importer of secologanin. To confirm and characterize this proposed role, we tested substrate range, transport directionality, and rate using the *X. laevis* expression system. *In vitro* transcription of *CrMATE1* was performed to produce cRNA for heterologous expression, where oocytes were injected with 25 ng of cRNA and incubated for 3 days to reach peak protein expression. By testing the directionality of substrate movement between oocytes (internal pH of 7) and the media at pH 5, this approach mimics *in planta* transport from the cytosol (pH 7) to the vacuole (pH 5). Thereby, export from the oocyte is a proxy for import into the vacuole and *vice versa*.

Substrate range assays were carried out with the secoiridoids, secologanin, loganic acid, and loganin. No vacuolar export was registered for any of the substrates (Figure 4A). Vacuolar import was tested by injecting the substrate into the oocyte, followed by assaying for 90 minutes in pH 5 media. No significant transport was detected for loganin and loganic acid (Figure 4B). Conversely, secologanin translocation was detected in both control (no *MATE*) and *MATE*-expressing samples. In the control oocytes, secologanin appears to partially diffuse or leak into the media; however, in *MATE* oocytes, all of the secologanin is successfully transported.

**Figure 4.**
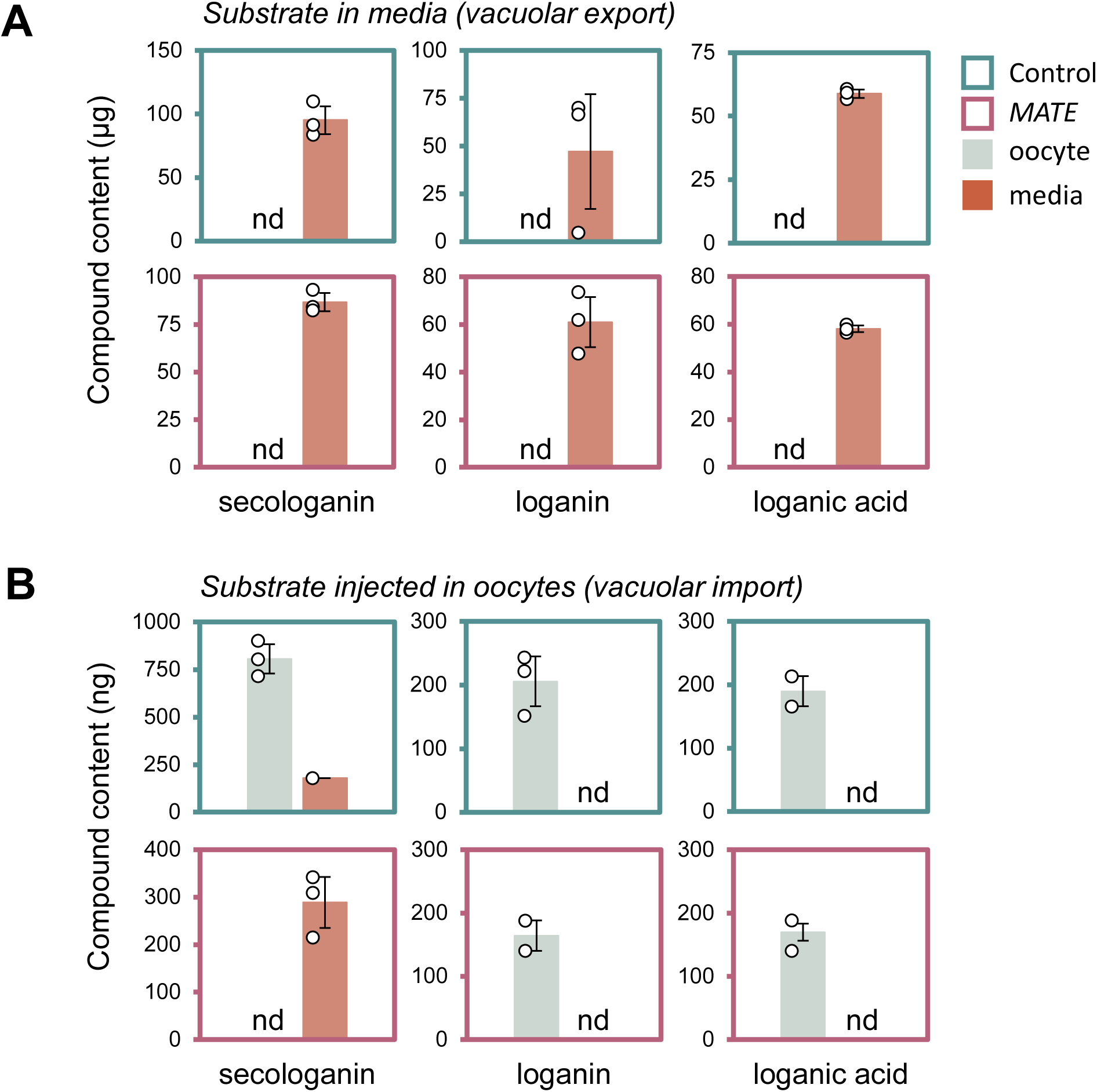
*In vivo* characterization of CrMATE1 for substrate range and transport directionality. Substrate specificity for **A)** vacuolar export was measured in control (non-injected) oocytes and oocytes expressing *CrMATE1*. Oocytes (3 x 5) were incubated in buffer supplemented with 100 µM of the secoiridoids, including secologanin, loganin, and loganic acid. **B)** Vacuolar import was assessed by injecting substrates into oocytes to a final internal concentration of 1 mM for secologanin and 100 µM for loganin and loganic acid. Each substrate was tested individually at pH 5 and assayed for 90 min. Substrate content in oocytes and media was determined by LC/Q-TOF MS. Data represent the mean ± S.D. of two or three biological replicates (data points shown).

A time-course assay (0-90 min, 7 time-points) further illustrated the rapid and complete translocation of secologanin into the media in *MATE*-expressing oocytes (25 min), compared to a slow and partial movement in control samples (Figures 5A, B). Strikingly, we noticed the accumulation of secologanol in control oocytes over time, mirroring the chemotyping results from *VIGS-MATE* leaves (Figure 5C). The rate of secologanin conversion to secologanol is faster than the passive diffusion or leakage of secologanin, where, at the end of a 90-minute incubation, there is 2-fold more secologanol in the oocyte than secologanin. On the other hand, no secologanol is detected within the oocytes of the *MATE*-expressing samples. Furthermore, no secologanol is detected in the media in either set of samples. The rapid transport of secologanin by CrMATE1 confirms its role as a highly efficient and substrate-specific vacuolar importer.

**Figure 5.**
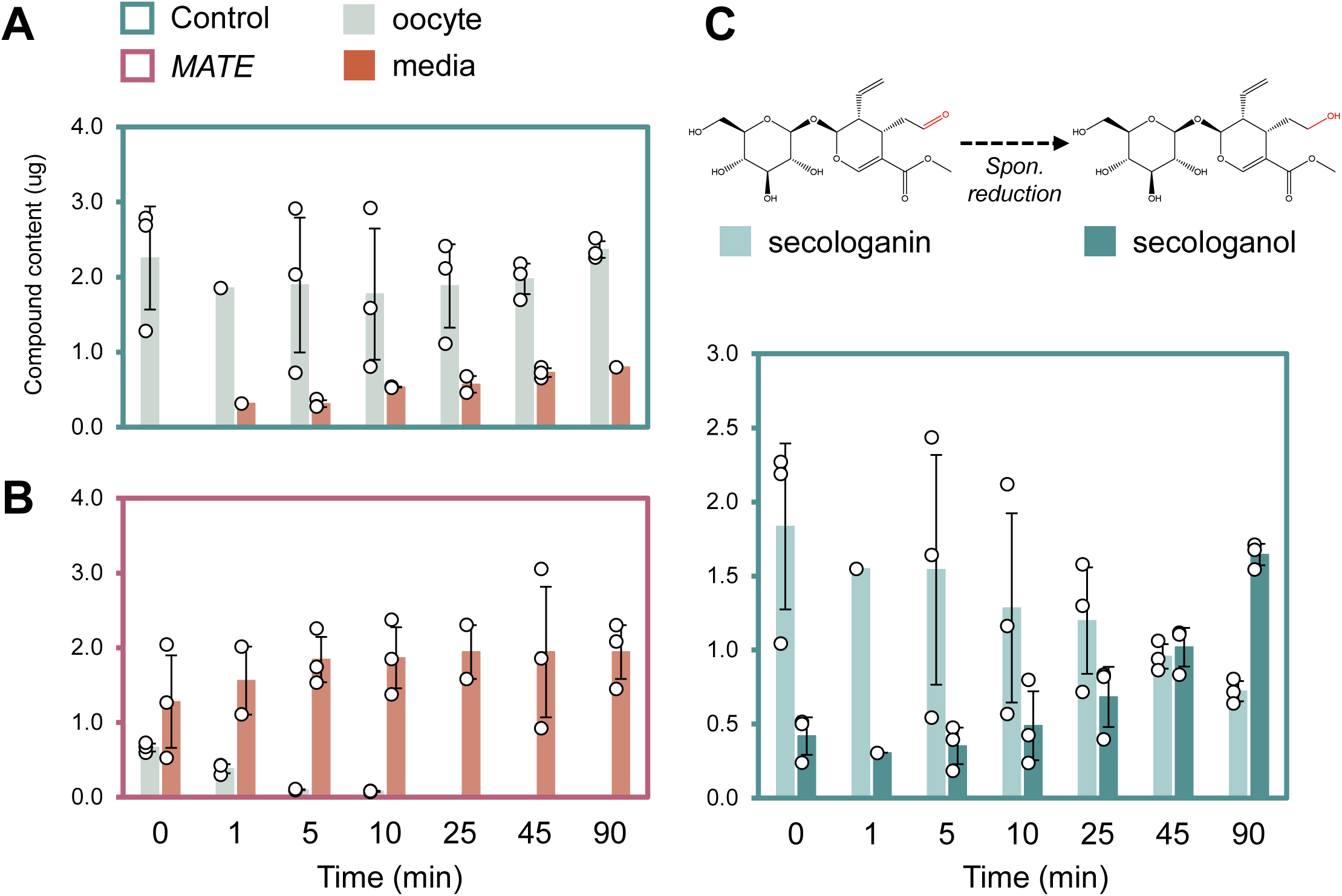
Rate of secologanin transport. **A)** Secologanin transport over time in control (non-injected) oocytes and **B)** *MATE*-expressing oocytes. Oocytes (3 x 5) were injected with the substrate to a final internal concentration of 1 mM and incubated for indicated times from 0-90 min (pH 5). Substrate content in oocytes and media was determined by LC/Q-TOF MS. **C)** The spontaneous reduction of secologanin was detected in non-injected oocytes, shown by the schematic legend indicating the position where the reduction is occurring. Amounts of secologanol (light turquoise) are compared to secologanin (turquoise) at each time-point.

## 3. DISCUSSION

Monoterpene indole alkaloid (MIA) biosynthesis in *C. roseus* is a complex pathway comprised of over thirty reactions that are highly compartmentalized, spanning multiple organelles and cell types (Figure 1) (St-Pierre *et al*., 1999; Courdavault *et al*., 2014; Carqueijeiro *et al*., 2016; Kulagina *et al*., 2022). Despite our current, intricate level of understanding, gaps remain in the description of inter- and intracellular transport of intermediates. Previously, three transport steps have been characterized at the plasma membrane and tonoplast of epidermal cells, executed by NPF transporters (CrNPF2.4-2.6 and 2.9) (Larsen *et al*., 2017; Payne *et al*., 2017) and an ABC transporter, CrTPT2 (Yu and De Luca, 2013). Here, we have fully characterized a *C. roseus* MATE transporter, CrMATE1, with a role in the vacuolar import of secologanin.

Referred to as *SLTr* by Li and colleagues (2023), they showed that silencing of *CrMATE1 in planta* only increased the levels of secologanol; secologanin and reported MIAs were not significantly changed. Our group has performed silencing in two varieties of *C. roseus* reporting consistent and drastic accumulation of secologanol, and a moderate but significant reduction of secologanin and the downstream MIA, catharanthine (Figures 2B, D). Reduced concentration of another MIA, vindoline, was variety-dependent. This chemical phenotype is likely due to the strong suppression of *CrMATE1* (73-80%) in the selected varieties (LD and PW) and under our conditions. The remainder of *CrMATE1* transcripts might function to ensure adequate transport of secologanin for the subsequent vacuolar condensation with tryptamine, forming strictosidine. On the other hand, we cannot rule out the possibility of other vacuolar importers of secologanin; however, the large build-up of secologanol suggests that CrMATE1 is of primary importance.

The substantial accumulation of secologanol in *MATE*-silenced plants appears to be a protective measure, reducing the cytotoxic aldehyde on secologanin, as the compound builds up due to lack of transport. The injection of secologanin into *X. laevis* control oocytes (without *CrMATE1*) resulted in a similar gradual build-up of secologanol (Figure 5C). We can rule out the spontaneous reduction of secologanin (at pH 5 or 7), as it is only detected within the control oocytes or in our VIGS plants. Different catalysts or factors might be responsible for formation of secologanol in either setting. On the other hand, *MATE*-expressing oocytes rapidly and completely translocate secologanin into the media, where it is stable and unchanged.

Another issue was the apparent leakage into the media of secologanin in control oocytes (Figures 4B, 5A). This problem appeared unique to secologanin, and we did not detect any passive diffusion of the accumulating secologanol (Figure 5). Therefore, it is unlikely that secologanin is passively diffusing; instead, it is likely leaking at the site of its injection or is carried out with the needle. The level of MATE-driven substrate translocation is significantly faster, and all the injected secologanin (1 mM) was moved into the media after 25 min (Figure 5B).

Finally, we confirmed the subcellular localization of CrMATE1 to the tonoplast with transient expression in *N. benthamiana* epidermal cells (Figure 3), consistent with its role as a vacuolar transporter. Meanwhile, the cognate transcript of *CrMATE1* is ubiquitously present throughout sampled tissues but with slightly higher expression in leaf tissues, where we expect a correlated higher degree of protein activity (Suppl. Figure 5). Indeed, leaf vacuoles are major sinks of alkaloid accumulation, and their translocation across the tonoplast for storage has also been characterized in *N. tabacum* MATE transporters NtMATE1/2 and NtJAT1/2 (Morita *et al*., 2009; Shoji *et al*., 2009; Shitan *et al*., 2014). In the case of MIA biosynthesis, CrMATE1 plays a role in controlling metabolic flux by delivering secologanin to its committing condensation reaction in the vacuole. Payne *et al*. (2017) show that the disruption of subsequent strictosidine export from the vacuole leads to cytotoxicity. Therefore, transvacuolar movement for both substrates and product can be an effective gatekeeping mechanism for this vital, yet physiologically harmful, compound.

Although leaves are the primary source of MIAs in *C. roseus*, biosynthesis also occurs in other organs like stems, flowers, and roots. Interestingly, secologanin and the preceding secoiridoids are mobile intermediates, moving from WT roots to low-secoiridoid mutant shoots in a grafted scion (Kidd *et al*., 2019). Therefore, the secologanin pool for leaf-MIA metabolism might not be sourced *de novo*, but can accumulate from inter-organ transport, potentially mobilized by another transporter(s) in conjunction with the epidermal importers, CrNPF2.4-2.6. Furthermore, the shuttling of secologanin into root-localized pathways, producing such compounds as vincadifformine, hörhammericine, and tabersonine-derivatives might require further unique transport proteins (Carqueijeiro *et al*., 2018; Williams *et al*., 2019). Overall, there remains salient gaps in our comprehension of inter-organelle and inter-cellular transport in MIA biosynthesis, including the vacuolar import of the other constituent for strictosidine formation, i.e., tryptamine.

The evolution of a dedicated transporter for secologanin is fascinating. We could speculate that it has neo-functionalized from a generic transporter of MEP pathway intermediates; however, there is no evidence for this theory. A phylogenetic analysis of MATE family amino acid sequences shows tonoplasts importers (CrMATE1, NtMATE1, NtMATE2, NtJAT2) clustering in *Group A* (Suppl. Figure 1), sharing several hydrophobic substitutions in their V-shaped pockets (Suppl. Figure 2). While a general V-shape structure is conserved among various MATEs, the nature of the pocket, including charge, polarity, and hydrophobicity, is highly variable. The crystal structure of CasMATE from the plant *Camelina sativa* has shed light on the role of critical residues within this pocket that determine substrate selectivity (Tanaka *et al*., 2021). The overall increased hydrophobicity of MATE proteins in group A is consistent with their role in transporting specialized metabolites with bulky aromatic rings.

Transporters are crucial in the matrix of specialized metabolism, mediating metabolic flux and anti-herbivory potency in balance with the homeostatic parameters of plant cells (Watkins and Facchini, 2022). The complete enzymatic resolution of the MIA pathway has made it possible for reconstitution into microbial and non-native plant systems, where recent efforts have made tremendous progress in the heterologous production of valuable MIAs (Zhang *et al*., 2022; Dudley *et al*., 2022; Shahsavarani *et al*., 2023; Gao *et al*., 2023). However, biosynthesis outside of the native context exposes many shortfalls, including liability for by-product formation and a loss of carbon to aberrant reactions (Dastmalchi *et al*., 2019a; Dastmalchi, 2021). Additionally, retrieving intermediates or substrates from the media can improve product titers. The utility of transporters was demonstrated by the inclusion of benzylisoquinoline alkaloid (BIA) uptake permeases (BUPs) in engineered yeast strains producing morphine and related intermediates (Dastmalchi *et al*., 2019b). The inclusion of BUPs in the co-cultured modules of the pathway increased titres up to 300-fold, allowing separation of the metabolic burden into three strains. Whether *C. roseus* transporters can improve heterologous production of MIAs has yet to be examined; however, it could be a notable consideration in providing an alternative supply chain for the valuable and life-saving compounds offered by this unassuming plant.

## 4. EXPERIMENTAL PROCEDURES

### Ethics Statement

All work involving animals was carried out under the McGill University Animal Use Protocol 2022-7758, authorized by the Animal Care Committee of the Office of Research Ethics and Compliance, McGill University.

### 4.1 Virus-Induced Gene Silencing

Virus-Induced Gene Silencing (VIGS) was performed, as previously described (Eng *et al*., 2022), using *C. roseus* var. “LD” and “PW” seedlings grown in a controlled chamber at 25 °C (16/8 h photoperiod). *Agrobacterium tumefaciens* (strain GV3101) cells either harbouring *pTRV2-MATE* (*VIGS-MATE*), *pTRV2-empty vector* (*EV*), or *pTRV2-CrPDS* (phytoene desaturase) were grown overnight (28 °C) and harvested by centrifugation. Cells were resuspended in infiltration buffer (10 mM MES pH 5.6, 10 mM MgCl_2_, 0.2 mM acetosyringone) to an OD_600_ of 1.5 and were cultured at 28 °C for 2.5 h. Cell suspensions of *pTRV1* and *pTRV2* were mixed, and a syringe needle was dipped in the mixture and used to penetrate the 4-week-old *C. roseus* seedlings underneath the shoot apical meristem. After penetration, an additional 0.12 mL of suspension was used to flood the wounded area. Thus, we produced plants co-transformed with *pTRV1* and *EV* (6 in LD, 6 in PW) or *VIGS-MATE* (6 LD, 9 PW) constructs. The infected seedlings were grown at a lower temperature of 20 °C until the *pTRV2-CrPDS* control seedlings started to show strong leaf bleaching (approximately 4 weeks post-injection). Silenced leaf pairs were harvested, split along the main vertical vein, and stored by flash-freezing. One half was used for RNA isolation using Trizol® reagent (Thermo Fisher, USA) and qRT-PCR studies, as described previously (Eng *et al*., 2022). The other half was used to extract leaf alkaloids; filtered extracts (5 μL) were submitted to LC-MS/MS for MIA quantifications. Changes in chemical profiles were evaluated by two-tailed, unpaired Student T-test.

### 4.2 qPCR of VIGS tissue

Quantitative Real Time-PCR (qRT-PCR) was performed on an Agilent AriaMx RealTime PCR instrument using the SensiFAST SYBR No-ROX qPCR 2X master mix (FroggaBio, Canada) according to the manufacturer’s protocol. The settings for the qRT-PCR (10 μL, 5 ng total RNA) included 40 cycles at 95 °C for 10 s and 60 °C for 30 s. The standard 2^−ΔΔCT^ method was used to quantify gene expression levels normalized to the expression of the *C. roseus*, *60S ribosomal RNA* housekeeping gene, *Cr60SrRNA* (Eng *et al*., 2022). Gene expression changes were analyzed by a two-tailed, unpaired Student T-test from independent biological samples and the average of three technical replicates.

### 4.3 Subcellular localization

The full-length *CrMATE1* gene was cloned into the entry vector, *pDONR-Zeo*, followed by recombination into the destination vector *pSITE-2CA* (Chakrabarty *et al*., 2007), for N-terminal fusion with eGFP. Vectors were constructed using Gateway^®^ Cloning technology (Invitrogen, USA). All constructs were confirmed by sequencing, and the corresponding plasmids were transformed into chemically competent *A. tumefaciens* cells (GV3101). Overnight cultures were grown to an OD_600_ of 0.4-.0.8 and resuspended in infiltration buffer to an OD_600_ of 1.0 (10 mM MES/KOH solution pH 5.6, 10 mM MgCl_2_, and 0.15 mM acetosyringone). Agroinfiltration was performed using a syringe without a needle on the abaxial side of the leaf according to the protocol by Sparkes *et al*. (2006).

*Nicotiana benthamiana* plants were grown for six weeks in a controlled chamber at a 22/20 °C (16/8 h photoperiod) and light intensity of 150 μmol/m^2^/s. Plants were sampled three days after infiltration, and leaf epidermal samples were visualized with a Leica SP5 CLSM equipped with a Radius 405 nm laser (Leica Microsystems, Germany). GFP fluorescence was assayed with excitation at 488 nm and emission at 510 nm. RFP fluorescence was assayed with excitation at 558 nm and emission at 583 nm. Samples were viewed with a 63x water immersion objective (microscope slide with coverslip thickness 1.5; 0.17 mm).

### 4.4 *In vitro* RNA transcription

The full-length coding sequence of *CrMATE1* was cloned into the *pTD2* oocyte expression vector (Duguet *et al*., 2016) and confirmed by sequencing, containing the gene of interest flanked by 5′- and 3′-*X. laevis* β-globin UTRs and a 3′ poly-A tail, respectively. Primers (Suppl. Table 1) were used to amplify *CrMATE1* CDS, plus flanking sites, with Q5® High-Fidelity DNA Polymerase (New England Biolabs, USA) from the *pTD2-MATE* construct, followed by gel-purification, using the Monarch® DNA Gel Extraction Kit (New England Biolabs, USA). *In vitro* transcription of this template was performed using the mMESSAGE mMACHINE T7 kit (Invitrogen, USA), precipitated with lithium chloride, and dissolved in RNase-free water. The final concentration was adjusted to 500 ng/µL for injection purposes.

### 4.5 *Xenopus laevis* oocyte extraction and injection

*Xenopus laevis* individuals used in this study were maintained following the McGill University Animal Use Protocol 2022-7758. Adult female frogs (Xenopus1, USA) were housed in the Xenoplus Housing 603 System (Technoplast, Italy). Frogs were anesthetized in 0.15% MS-222, pH 7.3 (Sigma-Aldrich, USA), an incision was made on the side of the abdomen for oocyte extraction, and oocytes were prepared according to standard protocol (Noonan and Beech, 2023). Lobes were placed in a Ca^2+^-free OR2 solution (82 mM NaCl, 2 mM KCl, 1 mM MgCl_2_, 5 mM HEPES buffer, pH 7.3) and separated into individual oocytes using fine tweezers. A 90-min incubation at RT on a tube rotator was followed by supplementation with 10 mg/mL collagenase type II (Sigma-Aldrich, USA) in Ca^2^-free OR2 and, finally, stopped by washing in Ca^2^-free OR2. Defolliculated oocytes were transferred to ND96 solution (NaCl 96 mM, KCl 2 mM, CaCl_2_ 1.8 mM, MgCl_2_ 1 mM and HEPES 5 mM, pH 7.3), supplemented with 2.5 mM sodium pyruvate (Sigma-Aldrich, USA) (Noonan and Beech, 2023). For injection, glass capillaries were pulled into injection needles (World Precision Instruments, USA), and the tips were clipped using tweezers. Thus prepared, needles were first backfilled with oil, followed by cRNA aspiration. Oocytes were injected the same day as the surgery with 50.6 nL (25 ng per injected gene) of cRNA using a Nanoject injector (Drummond Scientific, USA). After injection, oocytes were incubated for 3 days at 18 °C before assaying.

### 4.6 *In vivo* characterization of CrMATE1

For transport assays, oocytes were pre-incubated for 5 min in Kulori buffer (90 mM NaCl, 1 mM KCl, 1 mM CaCl_2_, 1 mM MgCl_2_, 5 mM MES, pH 5) to ensure intracellular steady-state pH before transferral to Kulori containing one of the tested compounds (secologanin, loganin, and loganic acid). Assays testing oocyte import were conducted by incubating control (non-injected oocyte) and MATE-expressing oocytes in Kulori buffer containing 100 µM of individual substrates. Oocyte export assays were conducted by injecting the test substrate into the oocyte to reach a final internal concentration of 1 mM for secologanin and 100 µM for loganin and loganic acid, assuming an internal oocyte volume of 1 µL. A 10-minute recovery period following substrate injection in ND96 (pH 7) was performed prior to transfer into Kulori buffer (pH 5). Assays were stopped after 90 min, unless stated otherwise, by washing the oocytes in 3 x 20 mL Kulori buffer without substrate. Media was collected and concentrated using a Savant SpeedVac DNA130 rotary-vacuum evaporator (Thermo Scientific, USA). Each assay was conducted with 3 biological replicates, each containing 5 oocytes. Pools of 5 oocytes were homogenized in 50 µL of 50% methanol. Homogenates were precipitated by centrifugation at 20,000 g for 10 min, frozen overnight, and followed by centrifugation at 20,000 g for 10 min, where supernatants of oocyte and media extracts were used for LC/QTOF-MS analyses. The presence and quantification of target compounds (substrates) within the oocytes and in the media were calculated using calibration curves of authentic standards. Data from the rate assay (Figure 5) was normalized with the dilution factor to calculate mass and corrected by a coefficient to remove technical error.

### 4.7 Liquid Chromatography-Mass Spectrometry

*In planta* samples from VIGS were measured with an Agilent Ultivo Triple Quadrupole LCMS (Agilent, USA) equipped with an Avantor® ACE® UltraCore™ SuperC18™ column (2.5 μm, 50 x 3 mm). For alkaloid detection from the variety “PW,” the following solvent system was used: solvent A, MeOH:ACN:1 M ammonium acetate:water at 29:71:2:398; solvent B, MeOH:ACN:1 M ammonium acetate:water at 130:320:0.25:49.7. The following linear gradient (8 min, 0.6 mL/min) was used: 0 min: 80% A, 20% B; 0.5 min: 80% A, 20% B; 5.5 min: 1% A, 99% B; 5.8 min: 1% A, 99% B; 6.5 min: 80% A, 20% B; 8 min: 80% A, 20% B. The solvent system for alkaloids extracted from the variety “LD” was as follows: solvent A, MeOH:ACN:5 mM ammonium acetate at 6:14:80; solvent B, MeOH:ACN:5 mM ammonium acetate at 24:64:10. The following linear elution gradient was used: 0-0.5 min: 99% A, 1% B at 0.3 mL/min; 0.5-0.6 min: 99% A, 1% B at 0.4 mL/min; 0.6-8.0 min: 1% A, 99% B at 0.4 mL/min; 8.0-8.3 min: 99% A, 1% B at 0.4 mL/min; 8.3-10.0 min: 99% A, 1% B at 0.3 mL/min. The photodiode array detector recorded from 200 to 500 nm. The MS/MS was operated with gas temperature at 300 °C, gas flow of 10 L/min, capillary voltage 4 kV, fragmentor 135 V, and collision energy 30 V in both polarities.

The Qualitative Analysis 10.0 software by Agilent was used for all LC-MS/MS analyses. The analytes were either dissolved in methanol or methanol: water in an equal volume ratio. Compounds were identified and quantified using peak areas (UV 280 nm: catharanthine; UV 300 nm: vindoline) using calibration curves for authentic standards. Secologanol was produced by reducing 150 μg secologanin with trace amounts of sodium borohydride dissolved in 150 μL methanol at room temperature for 1 h, which resulted in quantitative conversion. Secologanin and secologanol in VIGS experiments were quantified by standard curves of respective chemical standards recorded using MRM m/z 411 to 249 and m/z 413 to 251 ion transitions. Target MIA compounds were calculated per fresh sample weight.

Samples from *X. laevis* oocyte assays were measured using an Agilent 1290 Infinity II LC system coupled to the 6545 Q-TOF MS (Agilent Technologies, USA). The LC separation was conducted on a Poroshell 120 EC-C18 analytical column (Agilent, 2.1 × 5 mm, 1.8 μm). Samples (1 μL) were injected into the column, running on the mobile phases A (0.1% formic acid in water, 5 mM ammonium acetate) and B (MeOH:ACN 50:50 vol, 0.1% formic acid, 5 mM ammonium acetate). The LC gradient used was 0-0.5 min: 10% B; 0.5-2 min: ramp to 25% B; 2-4 min: ramp to 40% B; 4-6 min: ramp to 100% B; 6-8 min: 100% B; 8.5 min: 10% B; 8.5-10 min: 10% B at a flow rate of 0.4 mL/min. For targeted analysis, samples were run in negative ion mode: capillary voltage 3500 V, fragmentor voltage 175 V; skimmer 65 V; sheath gas temperature 350^°^C; sheath gas flow 12 L/min; Nebulizer 45 psig; scan rate 2 spectra/s; and a mass range of 50-1100 m/z. The MassHunter Quantitative Analysis software by Agilent was used for all LC/Q-TOF MS analyses.

## ACCESSION NUMBERS

*CrMATE1*: CRO_T006097/CRO_03G032350.1/AQM73450.1

## ACKNOWLEDGEMENTS

We are grateful to Dr. Robert Mullen (University of Guelph) for providing access to the Molecular and Cellular Imaging Facility and Dr. Satinder Gidda for direction and training. We also want to thank Reilly Pidgeon for his advice and Dr. Bastien Castagner for the use of their LC-MS, which was used for the initial analysis of samples. We thank Ball Seed^®^ (Illinois, USA) for supplying LD and PW seeds to Dr. Vincenzo De Luca for research.

## AUTHOR CONTRIBUTIONS

F.L. performed *in vivo* characterization and subsequent LC/Q-TOF MS data analyses. M.S. performed VIGS and subsequent LC-MS/MS analyses. Y.Q. supervised VIGS work and advised on analysis of results. C-J.H-H., V.M, and R.B. provided training and advice for *X. laevis* expression system. L.L. performed LC/Q-TOF, supervised by S.B. M.D. performed subcellular localization, phylogenetic analyses, and qPCR expression. The work began under the guidance of V.D.L. at Brock University and continued with his support. F.L. and M.D. wrote the manuscript with input from all co-authors.

## SHORT LEGENDS FOR SUPPORTING INFORMATION

**Supplementary Figure 1.** MATE proteins form six clades (groups A-F).

**Supplementary Figure 2.** Amino acid alignment of MATE family proteins.

**Supplementary Figure 3.** Suppression of *CrMATE1* transcript levels using virus-induced gene silencing.

**Supplementary Figure 4.** Subcellular localization of CrMATE1 co-expressed with a vacuolar marker.

**Supplementary Figure 5.** Relative expression of *CrMATE1*.

**Supplementary Table 1.** Nucleotide sequences of oligos used in this study.

