## Supplementary Materials for "*In vivo* characterization of a secologanin transporter from *Catharanthus roseus*"

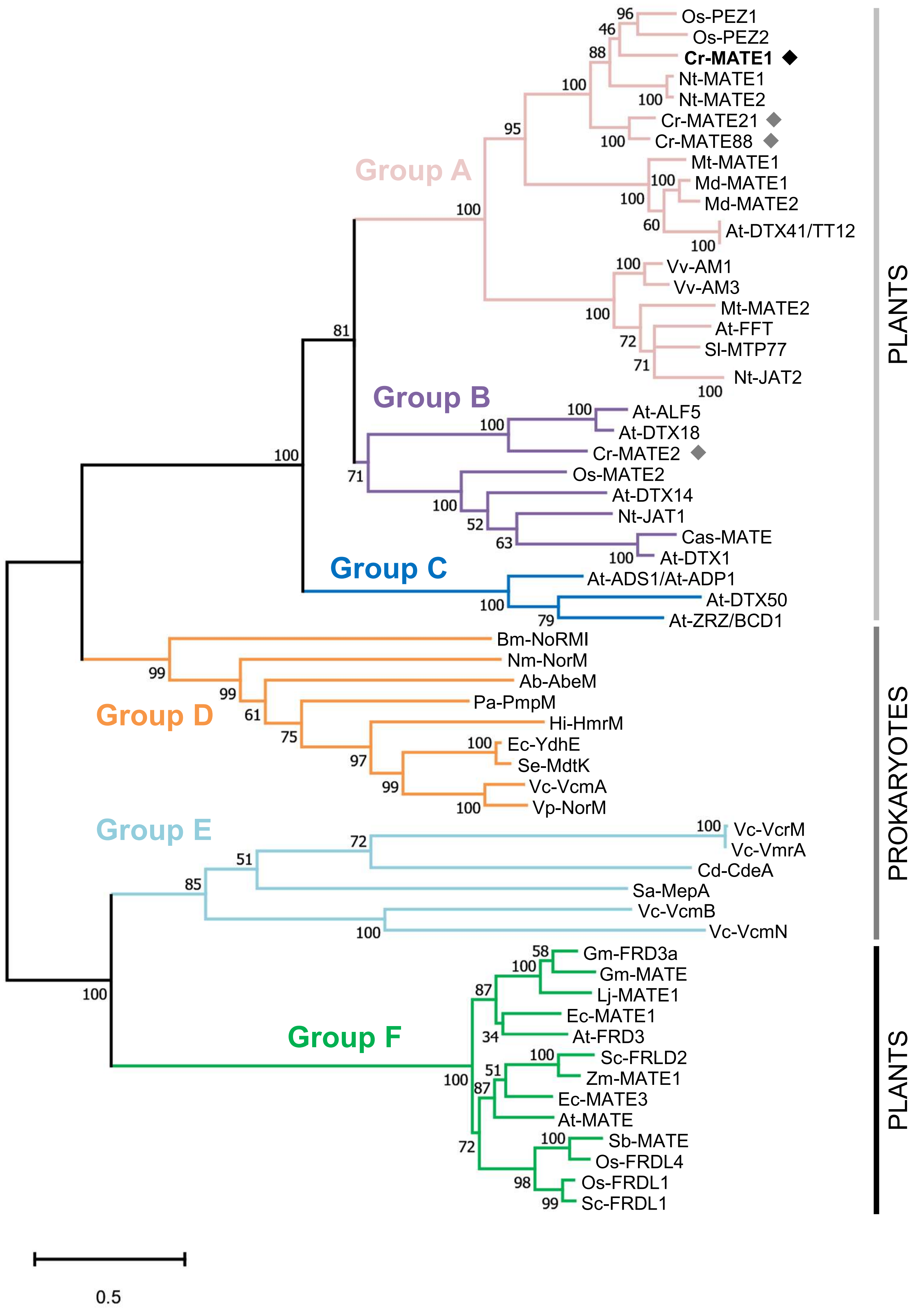


**Suppl. Figure 1**

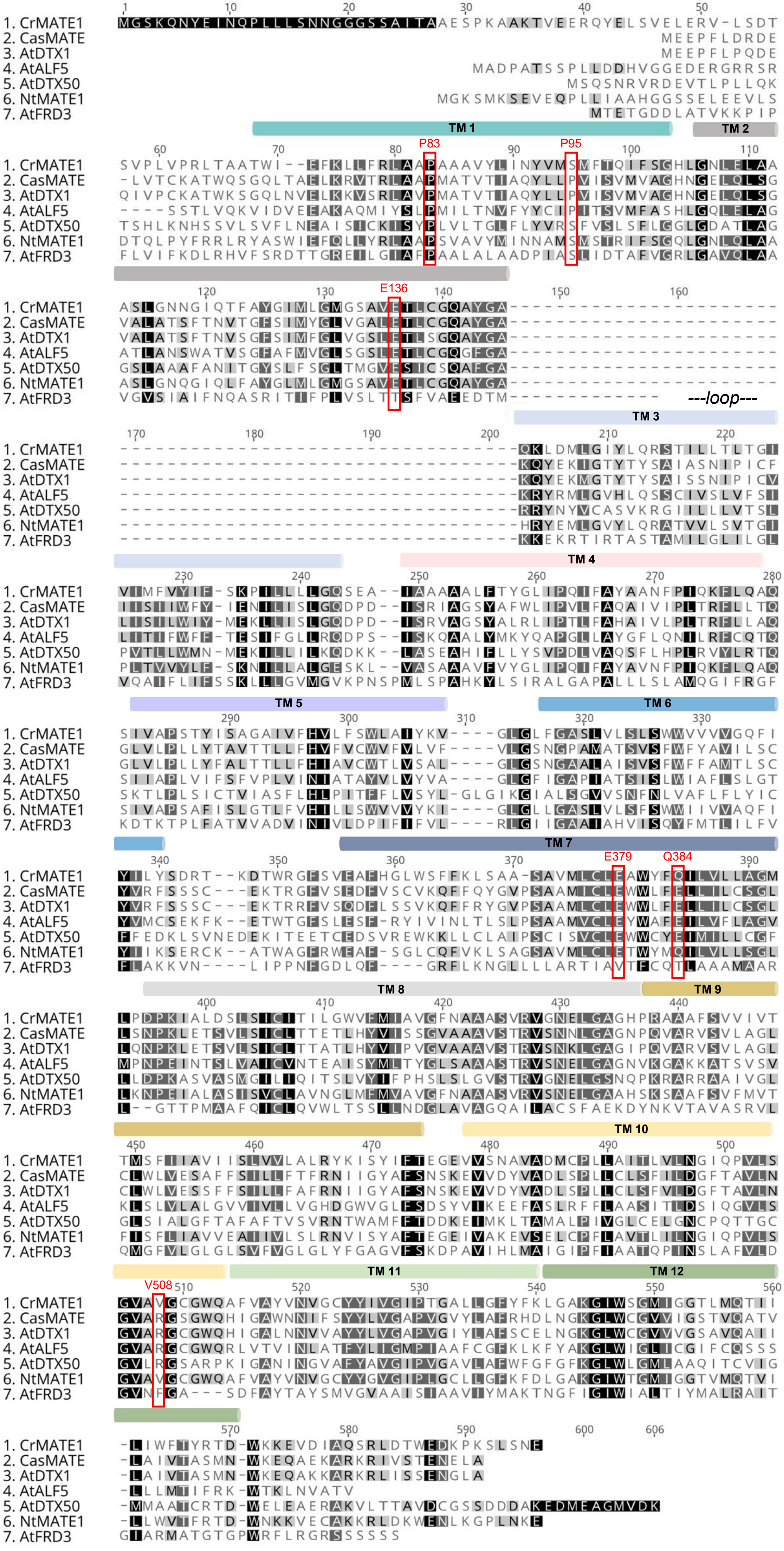


Suppl. Figure 2

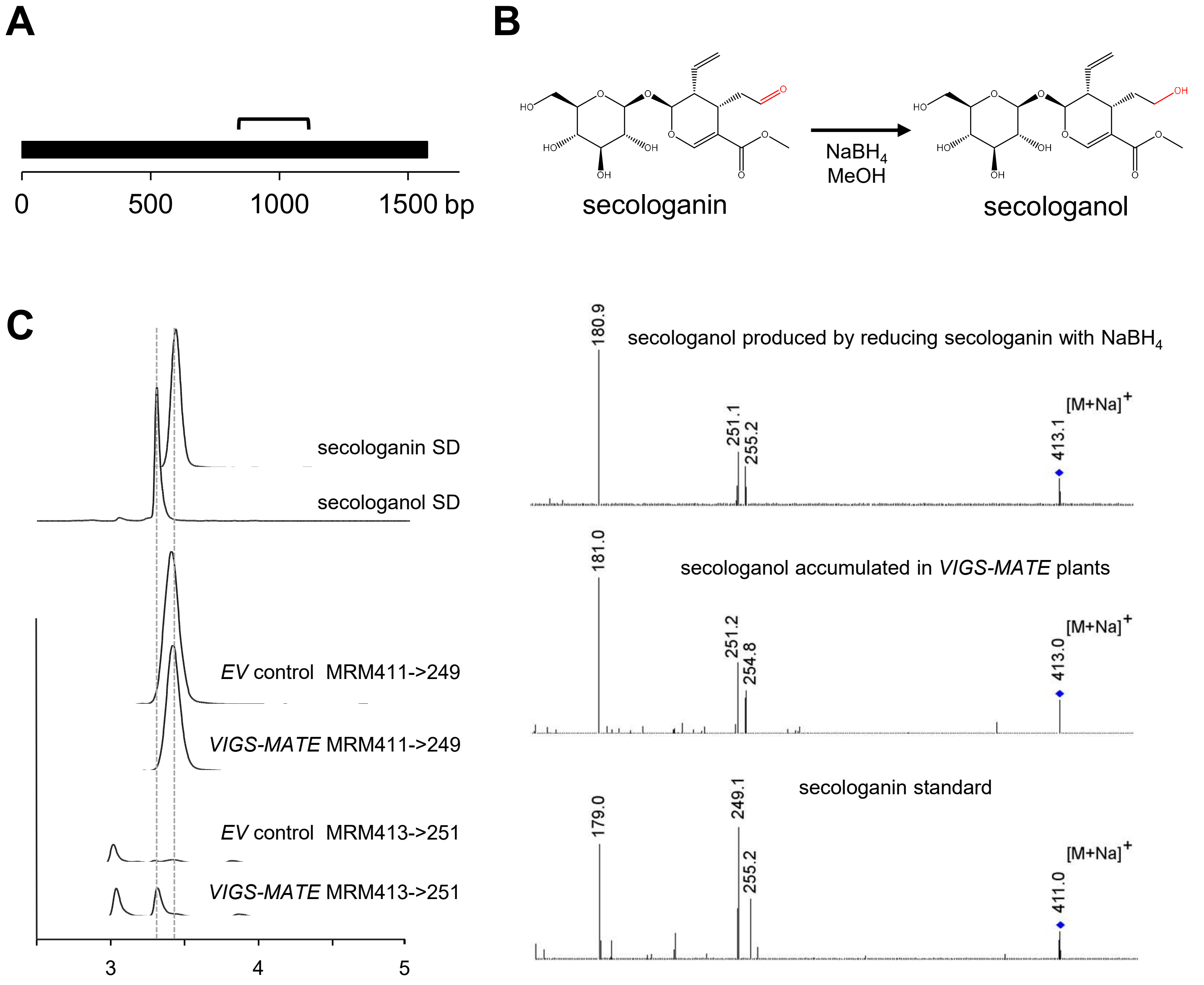


**Suppl. Figure 3**

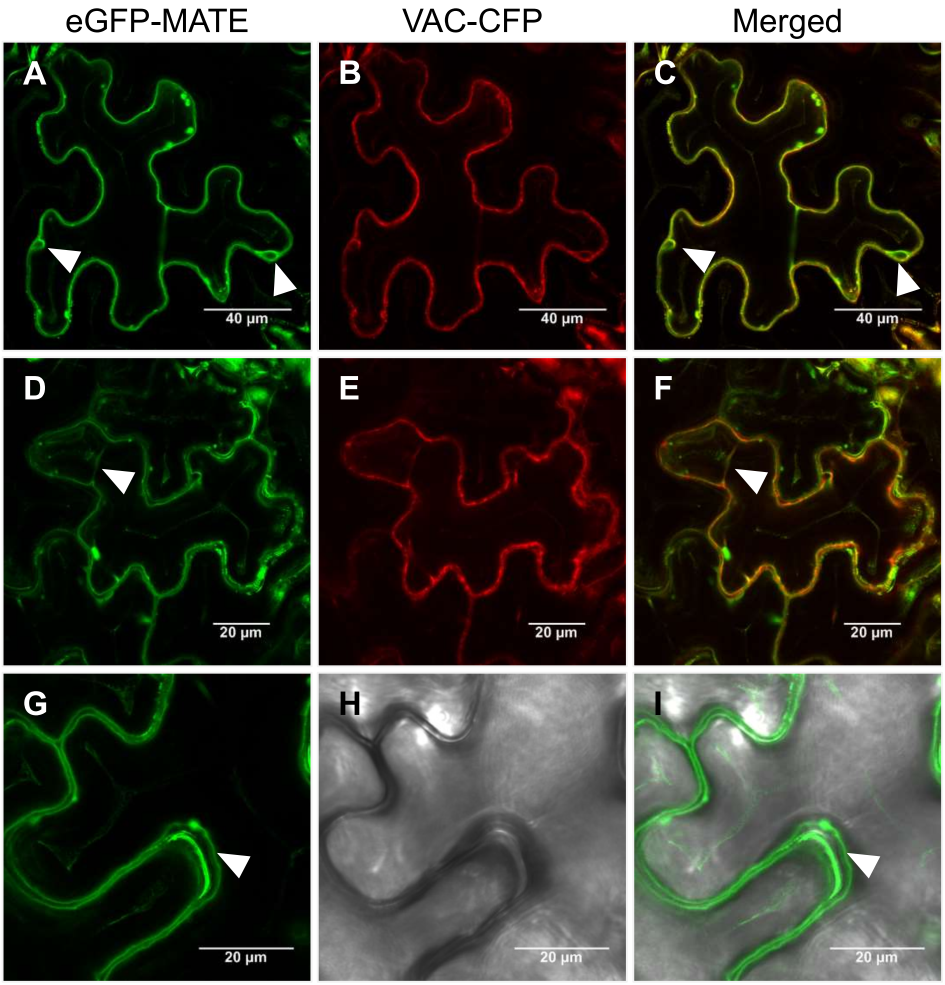


**Suppl. Figure 4**

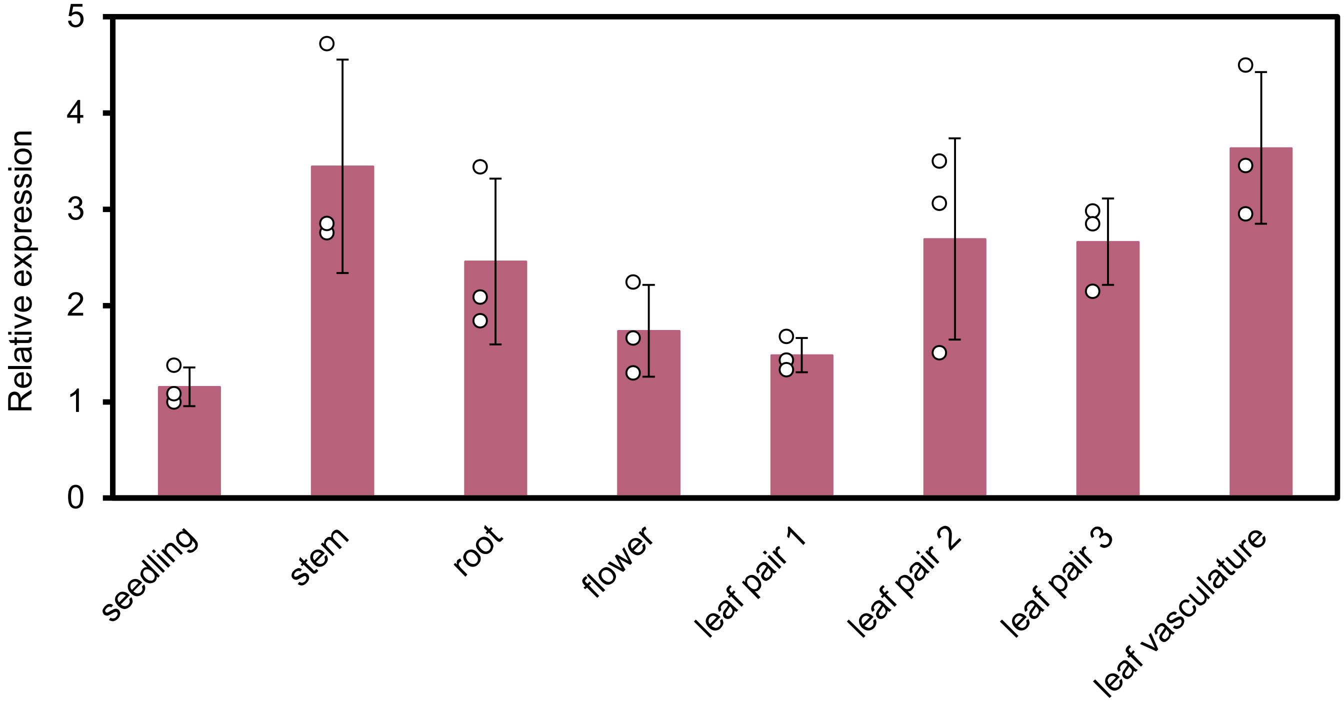


**Suppl. Figure 5**

| Primer name | Sequence (5' → 3') | Usage |
|---|---|---|
| VIGS-MATE-F | TTCGAATTCTTGTTCTTTTGGCTGGA | VIGS constructs |
| VIGS-MATE-R | GGTTGAATTCCATTAAGAACAAGA | VIGS constructs |
| qPCR-MATE-F | GACTTGGATTGTTCGGGGCATCT | qPCR of CrMATE1 |
| qPCR-MATE-R | AATCCACGCCAAGTGTCCTTAGTC | qPCR of CrMATE1 |
| qRT-PCR-60S-F | TCTTAGTTGGAATGTTCAGCACCTG | qPCR reference gene |
| qRT-PCR-60S-R | CAAGGTTGGAGCCCCTGCTCGTGTT | qPCR reference gene |
| CrMATE104-FWD | GGGGACAAGTTTGTACAAAAAAGCAGGCTTTATGGGTTCCAAACAAAACT | subcell. localization |
| CrMATE104-REV | GGGGACCACTTTGTACAAGAAAGCTGGGTTTATTCATTGGACAAAGATTTTGG | subcell. localization |
| CrMATE101-FWD | GGGGACAAGTTTGTACAAAAAAGCAGGCTATGGGTTCCAAACAAAACT | subcell. localization |
| CrMATE101-REV | GGGGACCACTTTGTACAAGAAAGCTGGGTATTCATTGGACAAAGATTTTGG | subcell. localization |
| MATE-not1-fwd | GAAATATGCGGCCGCATGGGTTCCAAACAAAAC | cloning CrMATE1 into pTD2 |
| MATE-apa1-rev | CTTTATAGGGCCCTTATTCATTGGACAAAGATTTTGGC | cloning CrMATE1 into pTD2 |
| pTD2_F | TTGGCACCAAAATCAACGGG | linearizing template for IVT of CrMATE1 |
| SP6_R | GATTACGCCAAGCTATTTAGGTGACAC | linearizing template for IVT of CrMATE1 |
| **Additional sequences** | **(5' → 3')** | |
| CrMATE1 VIGS portion | ATTCTTGTTCTTTTGGCTGGAATGCTTCCTGATCCTAAAATCGCTTTGGATTCCCT<br>CTCCATTTGCATTACAATCTTGGGTTGGGTATTCATGATAGCCGTTGGATTCAAT<br>GCTGCTGCCAGTGTGAGAGTAGGGAATGAACTAGGGGCAGGACATCCAAGGGC<br>AGCTGCATTTTCAGTAGTAATAGTGACAACAATGTCATTCATAATAGCAGTGATAA<br>TATCATTAGTGGTACTTGCTTTGCGCTTCAAAATTAGCTATATCTTTACCGAAGGT<br>GAAGTTGTAAGCAATGCTGTTGCCGATATGTGTCCCTTGCTCGCCATCACTCTTG<br>TTCTTAATGGAATTC | |

**Suppl. Table 1**
